# Lenacapavir-induced capsid damage exposes HIV-1 genomes from nuclear speckles

**DOI:** 10.1101/2025.07.07.663551

**Authors:** Thorsten G. Müller, Severina Klaus, Vojtech Zila, Svenja L. Nopper, Gonen Golani, Vera Sonntag-Buck, Anke-Mareil Heuser, Maria Anders-Össwein, Ulrich Schwarz, Vibor Laketa, Barbara Müller, Hans-Georg Kräusslich

## Abstract

Following cell entry, HIV-1 capsids enter the nucleus by passage through nuclear pores and reach nuclear speckles, where uncoating of the reverse-transcribed genome and integration occur. Here, we characterized the ultrastructure of HIV-1 subviral complexes in nuclei of primary monocyte-derived macrophages and cell lines using live-cell imaging, super-resolution microscopy and correlative light and electron tomography in the absence and presence of capsid-targeting inhibitors Lenacapavir and PF74. Capsid-like structures containing viral DNA as well as broken capsids clustered in nuclear speckles and were displaced from speckles by drug treatment. This was accompanied by alteration of the nuclear capsid structure, with electron-dense protrusions emanating from the narrow end of capsid cones and exposure of complete, or almost complete, genomic HIV-1 DNA. Our data indicate that synthesis of genomic dsDNA can be completed inside the closed HIV-1 capsid, and speckle-associated factors appear to be required to promote genome uncoating. This may ensure that genome uncoating occurs at optimal sites for integration into transcriptionally active chromatin. The results also shed further light on the mechanism of action of Lenacapavir.

## Introduction

Human immunodeficiency virus type 1 (HIV-1) replicates in CD4^+^ T cells and non-dividing macrophages of infected individuals. Within the last three decades, pharmacologically inhibiting viral enzymes has transformed a deadly infection into a life-long chronic condition, yet several challenges, including resistant mutations, drug adherence, and availability issues have prevented global elimination. The recent discovery, that the viral structural capsid protein (CA) shell plays a central role for the intertwined processes essential for replication (reviewed in (Müller *et al*, 2022; Jang & Engelman, 2023; Morling *et al*, 2025)) accelerated tackling some of these challenges with a novel class of drugs (Blair *et al*, 2010; Bester *et al*, 2020; Link *et al*, 2020) called capsid inhibitors. One of these capsid inhibitors termed Lenacapavir (LEN, GS-6207, or brand name “Sunlenca”) (Bester *et al*, 2020; Link *et al*, 2020) was recently approved by the European Medicines Agency and U.S. Food and Drug Administration for use in patients with multi-drug resistant HIV infections (Paik, 2022), and shows unprecedented efficacy in preexposure prophylaxis (Bekker *et al*, 2024), culminating in the selection of LEN as the “2024 Breakthrough of the Year” by Science magazine (Cohen, 2024).

The conical HIV-1 capsid is composed of a lattice comprising ∼ 250 hexamers and 12 pentamers of CA, with its unique structure playing a key role for many of its functions (Sundquist & Kräusslich, 2012; Freed, 2015). Upon arrival in the host cell by fusion of the viral membrane with the plasma membrane, the capsid shell facilitates transport towards the nucleus and shields the viral genome from the innate immune system. Simultaneously, the capsid acts as reaction container for reverse transcription, converting the viral single stranded RNA genome into double stranded DNA (dsDNA). This process starts following entry of the capsid into the cytoplasm (Hu & Hughes, 2012), but is only completed inside the nucleus (Dharan *et al*, 2020; Selyutina *et al*, 2020; Burdick *et al*, 2020; Francis *et al*, 2020; Rensen *et al*, 2021; Müller *et al*, 2021). The cytoplasmic capsid encasing the active replication complex then enters the nucleus, where the viral dsDNA genome is released from the capsid in a process called uncoating (Dharan *et al*, 2020; Burdick *et al*, 2020; Selyutina *et al*, 2020; Müller *et al*, 2021, 2022; Zila *et al*, 2021; Gifford & Melikyan, 2024; Kreysing *et al*, 2025). The capsid integrity appears to stay mostly intact until shortly before genome integration (Li *et al*, 2021), which preferentially occurs into nuclear speckle-associated chromatin domains (Sowd *et al*, 2016; Achuthan *et al*, 2018; Francis *et al*, 2020), to ultimately drive expression of the viral proteins and ensure long-term persistence.

Nuclear entry requires transport of the viral genome and associated proteins through nuclear pore complexes (NPC). Recent evidence has shown that seemingly intact capsids enter the nucleus through nuclear pores in different cell types (Zila *et al*, 2021; Müller *et al*, 2021; Schifferdecker *et al*, 2022), including primary human monocyte-derived macrophages (MDM) (Kreysing *et al*, 2025). This requires interactions between a hydrophobic binding cleft within the capsid lattice and FG-nucleoporins (Nups) of the NPC (Fu *et al*, 2024; Xue *et al*, 2023; Dickson *et al*, 2024). Being the largest nuclear import cargo known to date, capsid nuclear entry represents a sterical challenge, which can induce damage to the NPC (Kreysing *et al*, 2025), with the elastic properties of its lattice structure proposed to contribute to maintaining capsid integrity (Deshpande *et al*, 2024). At the nucleoplasmic side of the NPC, the host factor cleavage and polyadenylation specificity factor subunit 6 (CPSF6) binds to the same binding cleft in the capsid lattice as FG-Nups and mediates capsid release from the NPC (Bejarano *et al*, 2019; Achuthan *et al*, 2018; Chin *et al*, 2015; Zila *et al*, 2019). Subsequently, capsids migrate to nuclear speckles (Francis *et al*, 2020; Li *et al*, 2020; Selyutina *et al*, 2020; Rohlfes *et al*, 2025). Clustering of incoming viral structures has been observed inside the nucleus of tissue-culture adapted cell lines (Rensen *et al*, 2021; Schifferdecker *et al*, 2022; Müller *et al*, 2021).

The molecular mechanisms driving uncoating and destabilisation of the capsid shell are still not completely understood. A recent report suggested that completion of reverse transcription resulting in production of full length viral cDNA induces uncoating (Burdick *et al*, 2024). The additional presence in the capsid and the relatively rigid nature of the genomic dsDNA (Garcia *et al*, 2007) – as opposed to the flexible RNA (Chen *et al*, 2012) – may exert a disruptive mechanical outward force on the capsid. Accordingly, genome length of HIV-1 based vectors correlated with transduction efficiency indicating a critical minimal genome length of ∼ 6 kb for efficient uncoating (Burdick *et al*, 2024). This model gained support from theoretical analyses (Rouzina & Bruinsma, 2014) and *in vitro* studies, in which isolated HIV-1 capsids were analysed by atomic force microscopy (Rankovic *et al*, 2017, 2018, 2021; Xu *et al*, 2020) or cryo-EM (Christensen *et al*, 2020) after undergoing endogenous reverse transcription *in vitro*. While these results clearly point to an important role of genome length for uncoating, it is currently not clear whether completion of reverse transcription of the HIV-1 genome is sufficient or whether additional uncoating factors are required.

LEN (Bester *et al*, 2020; Link *et al*, 2020) as well as the previously described compound PF74 (Blair *et al*, 2010) inhibit viral infection in the early phase with multiple modes of action. These compounds bind to the same hydrophobic cleft between adjacent capsid hexamers as FG-Nups and CPSF6 (Bhattacharya *et al*, 2014; Bester *et al*, 2020). Compound binding stabilizes the hexameric lattice, but is incompatible with CA pentamers (Huang *et al*, 2025; Bhattacharya *et al*, 2014; Márquez *et al*, 2018; Faysal *et al*, 2024). This is consistent with experimental findings that PF74 or LEN addition to capsids with an internal fluid phase marker led to capsid breakage and release of the marker *in vitro*, while most of the hexameric capsid lattice remained intact (Márquez *et al*, 2018; Faysal *et al*, 2024; Li *et al*, 2025). Drug binding also competitively inhibits capsid interaction with FG-repeats of Nups and CPSF6 (Bhattacharya *et al*, 2014; Bester *et al*, 2020), and therefore blocks nuclear entry and subsequent nucleoplasmic trafficking to nuclear speckles. These effects occur at much lower concentrations of LEN or PF74 than lattice disruption. Furthermore, PF74 has been shown to displace subviral HIV-1 complexes from nuclear speckles (Francis *et al*, 2020) and to reveal CA epitopes for immunostaining (Müller *et al*, 2021), both features dependent on CPSF6 coating of the capsid.

While much is known about drug binding and its effect on capsid structure and nuclear entry, their effects on subviral HIV-1 complexes inside the nucleus and the ultrastructure of the nuclear capsid are not well understood. Here, we analysed HIV-1 capsids and their retention within nuclear speckles in the absence or presence of LEN/PF74 in MDM and cell lines using correlative light and electron microscopy (CLEM), electron tomography (ET), and (super-resolution) fluorescence microscopy in combination with live-cell detection of viral DNA. For visualization of reverse transcribed viral genomes, we employed either labelling of newly synthesized DNA by the nucleoside analogue 5-ethynyl-2′-deoxyuridine (EdU) (Peng *et al*, 2015) or specific detection of HIV-1 dsDNA by the ANCHOR system (Saad *et al*, 2014). The latter approach is based on the prokaryotic chromosomal partitioning system ParB-*parS* and requires almost complete reverse transcription of the HIV-1 genome as well as accessibility of the viral dsDNA for detection (Müller *et al*, 2021).

We observed rapid exposure of previously hidden pre-synthesized nuclear HIV-1 genomic dsDNA upon LEN treatment of infected cells, which coincided with ultrastructural alterations including bifurcated protrusions at the narrow end of capsid cones displaced from nuclear speckles. These observations indicate that complete or almost complete viral dsDNA synthesis can occur within the CPSF6-coated nuclear HIV-1 capsid or a capsid-like structure without immediate capsid breakage, thereby suggesting that additional factors, besides dsDNA synthesis, may be needed to trigger uncoating.

## Results

### HIV-1 capsid structures cluster in nuclear speckles of primary monocyte-derived macrophages

We first aimed to characterize the fate of HIV-1 capsids beyond the nuclear pore in terminally differentiated primary human MDM. We have previously observed clustering of HIV-1 capsids in the nuclei of HeLa-derived cell lines (Müller *et al*, 2021; Schifferdecker *et al*, 2022), and in the SupT1 T-cell line (Zila *et al*, 2021; Schifferdecker *et al*, 2022). In MDM, HIV-1 clusters of cDNA and fluorescently labelled IN largely co-localizing with nuclear speckles have been reported by others (Rensen *et al*, 2021; Francis *et al*, 2020). In order to determine whether these genome clusters also correspond to accumulations of capsids or capsid-like structures in primary MDM, we employed super-resolution fluorescence microscopy and CLEM to characterize the ultrastructure of subviral complexes associated with nuclear speckles.

Analyses were performed using an HIV-1 NL4-3 derivative carrying a deletion in the regulatory protein Tat and point mutations in the integrase (IN) active site (NNHIV; (Zila *et al*, 2021; Müller *et al*, 2021)). This variant is non-replication competent, but undergoes all post entry-events before the integration stage. For detection of subviral HIV-1 complexes, we employed a fluorescent Vpr-IN fusion protein provided *in trans* during particle production (Albanese *et al*, 2008), resulting in incorporation of labeled IN into the capsid. In previous studies, labeled IN was found to remain associated with capsid-like structures in the cytosol and nucleus in various cell types (Zila *et al*, 2021; Müller *et al*, 2021), and stayed associated with structures resembling capsid remnants upon separation from the viral cDNA (Müller *et al*, 2021). Reverse transcribing HIV-1 replication complexes in infected cells were detected by incorporation of the nucleoside analog EdU into the nascent viral DNA, followed by click-labeling. Although EdU incorporation is non-specific, this strategy labels almost exclusively viral replication complexes inside the nuclei of terminally differentiated cells lacking cellular DNA synthesis (Peng *et al*, 2015; Bejarano *et al*, 2019; Stultz *et al*, 2017; Francis *et al*, 2020; Rensen *et al*, 2021; Müller *et al*, 2021). Nuclear speckles were identified by immunostaining with the antibody SC35, which recognizes the major speckle protein SRRM2 (Ilik *et al*, 2020).

MDM infected with the labeled virus were stained and nuclear subviral complexes were imaged at 72 h post infection (p.i.) by Airyscan confocal microscopy. This analysis revealed a clear colocalization of IN.eGFP and EdU punctae within SRRM2-positive compartments (Figure 1a, Supplemental Figure 1), consistent with previously published findings (Rensen *et al*, 2021; Francis *et al*, 2020). To obtain a more detailed view of CA distribution within nuclear speckles, we performed dual-color STED nanoscopy of infected cells immunostained against SRRM2 and HIV-1 CA. SRRM2 did not appear homogeneously distributed within the speckle region at nanoscopic resolution (<50 nm; Figure 1b), but formed distinct and separated subclusters in the range of ∼ 50-100 nm. STED images revealed multiple CA positive objects clustering in internal speckle regions between these SRRM2 subclusters (Figure 1b,c). Their size and labeling intensity indicated that these objects may not represent individual capsids, but rather correspond to accumulations of capsids and/or capsid remnants (Figure 1b,c).

**Figure 1.**
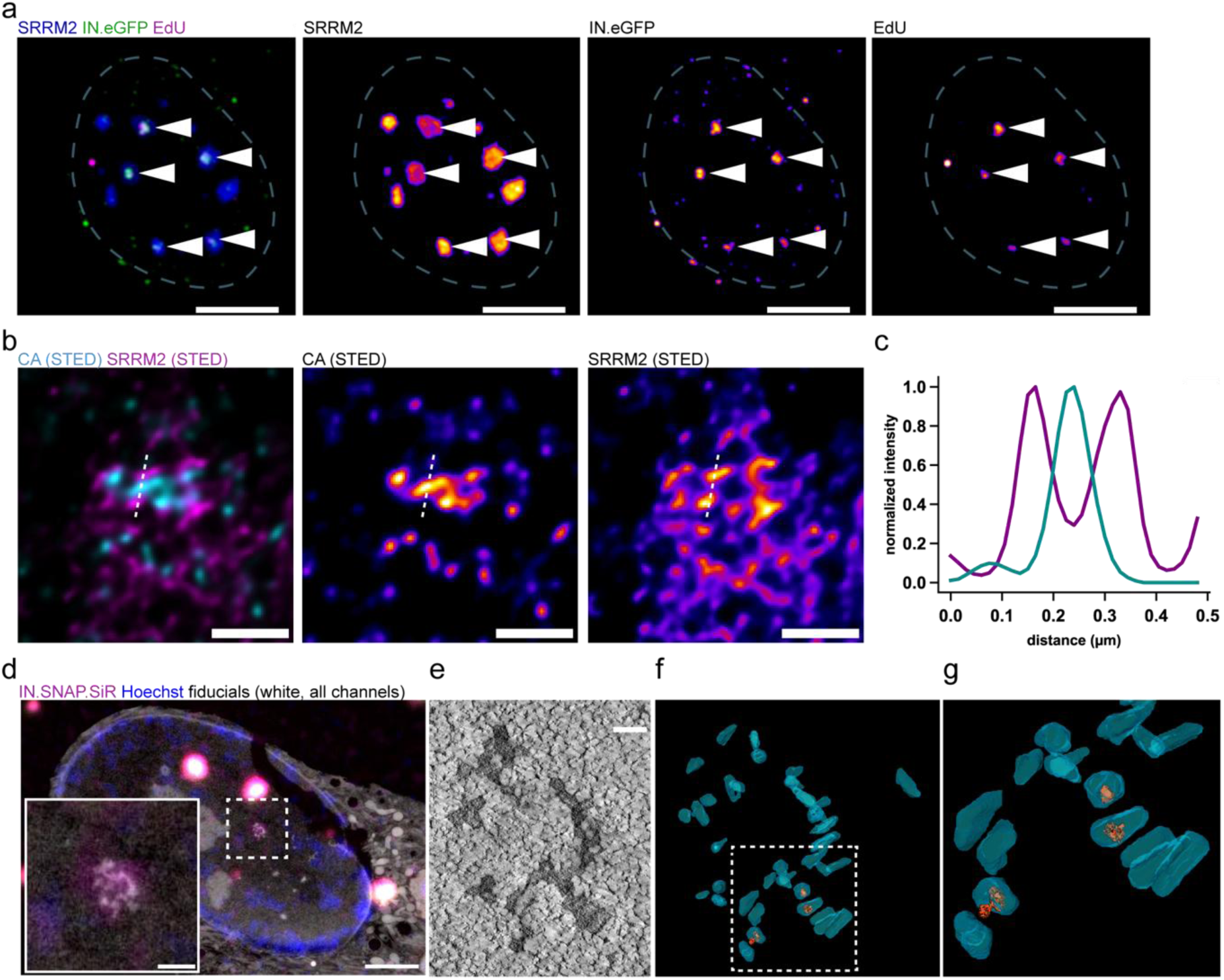
Super resolution and CLEM-ET analysis of capsid clusters in nuclear speckles of MDM. **a-c.** Super-resolution analysis of HIV-1 cDNA within nuclear speckles of MDM showing EdU IN.eGFP (a) and CA (b) signals in a central region of SRRM2 condensates. **a.** Representative image of a maximum intensity projection of an MDM nucleus (white dashed line). Cells were infected for 72 h with VSV-G pseudotyped NNHIV carrying INeGFP (green) in presence of EdU followed by fixation, EdU click labeling (magenta) and immunofluorescence staining using an antibody against SRRM2 (blue). Samples were imaged using Airyscan microscopy. Scale bar: 5 µm **b.** Dual color STED images of immunostained CA and SRRM2 of MDM infected with VSV-G pseudotyped NNHIV for 72 h. Scale bars: 0.5 µm **c.** Quantification of signal intensities along the white dotted line in (b) normalized to the highest value. **d-g.** CLEM-ET analysis of infected MDM. Cells were cryo-immobilized by high-pressure freezing followed by freeze-substitution and plastic embedding. **d.** CLEM overlay (with inverted EM image) of the 250 nm thin section of the cell, positive for IN.SNAP.SiR (magenta), post-stained with Hoechst (blue), and decorated with multifluorescent fiducials for correlation (all channels, white). Scale bars: 2.5 µm (overview) and 500 nm (enlargement). **e.** A single slice of the reconstructed electron tomogram correlated to the IN.SNAP.SiR signal boxed in (d.). Scale bar: 100 nm. **f,g.** 3D rendering of the tomogram shown in (e.) and enlargement of the boxed region in (f). See Supplementary Movie 1.

To further elucidate the ultrastructure of these viral objects, we applied CLEM-ET, using IN fused to the self-labelling SNAP-tag and stained with silicone rhodamine (SiR) as a marker to target the positions of nuclear subviral complexes. Infected MDM were subjected to cryo-immobilization by high-pressure freezing, followed by freeze-substitution and plastic embedding. Positions of interest were identified in thin sections based on IN.SNAP.SiR fluorescence using light microscopy and analyzed by ET. Clusters of elongated or conical electron dense objects with sizes consistent with HIV-1 capsids were observed at positions of IN.SNAP.SiR signals (Figure 1d-g). Some of these capsids appeared morphologically indistinguishable from virion-associated capsids, while broken structures and capsid remnants were also observed. Some structures contained internal density, likely representing the viral nucleic acid complex, while others appeared empty (Figure 1d-g). These data support recent findings that seemingly intact capsids with active reverse transcription can enter the nucleus of MDM (Kreysing *et al*, 2025) and reveal their subsequent accumulation and breakage in clusters within the central region of nuclear speckles.

### Nuclear speckle localization of capsids can be pharmacologically perturbed using LEN

Translocation of subviral HIV-1 complexes to nuclear speckles and their retention in the speckle area have been shown to be dependent on CPSF6, which forms a layer on the capsid inside the nucleus (Francis *et al*, 2020; Li *et al*, 2020; Selyutina *et al*, 2020; Rohlfes *et al*, 2025; Bejarano *et al*, 2019). Given the overlapping binding site of LEN and CPSF6, we analyzed the influence of LEN or PF74 on retention of speckle-associated HIV-1 capsids in MDM.

First, we determined the LEN concentration required for displacement of CPSF6 from nuclear capsids in MDM. To ensure completion of the post-entry phase, drug treatment was initiated at 72 h p.i. MDM infected with IN.eGFP labelled particles were treated with different concentrations of LEN for 1 h, immunostained for CPSF6 or CA and imaged using quantitative 3D confocal spinning disc microscopy. Full displacement of CPSF6 from nuclear subviral complexes in MDM was observed at a concentration of 500 nM LEN (Figure 2a,b). Concomitantly, CA epitopes were exposed on these complexes, enabling efficient immunostaining (Figure 2c,d). The same effect was achieved with 15 µM PF74 in primary MDM (Supplemental Figure 2a-d), similar to results obtained with PF74 in model cell lines and T cells (Müller *et al*, 2021; Ay *et al*, 2025). The LEN concentration required to displace CPSF6 from nuclear HIV-1 capsids was 10-fold lower in HeLa-based cells compared to MDM; 5 nM LEN and 50 nM LEN, respectively, resulted in partial and full displacement of CPSF6 and CA epitope exposure in HeLa-based cells (Supplemental Figure 3a-d). LEN binds the capsid hexameric lattice with a K_D_ of ∼ 200 pM (Faysal *et al*, 2024; Briganti *et al*, 2025) and the EC50 for inhibition of replication is at ∼ 50 pM in several cell lines. However, LEN inhibits different post-entry steps at different concentrations (Bester *et al*, 2020). Nuclear import of HIV-1 complexes is completely blocked at 5 nM LEN in Hela-based cells, while reverse transcription requires a concentration of 50 nM for full inhibition (Bester *et al*, 2020). Hence, the concentration of LEN required to fully displace CPSF6 within one hour of drug treatment of Hela-based cells is in a similar range as the concentration needed to block reverse transcription, but notably higher than the concentration required to block nuclear import in these cells.

**Figure 2.**
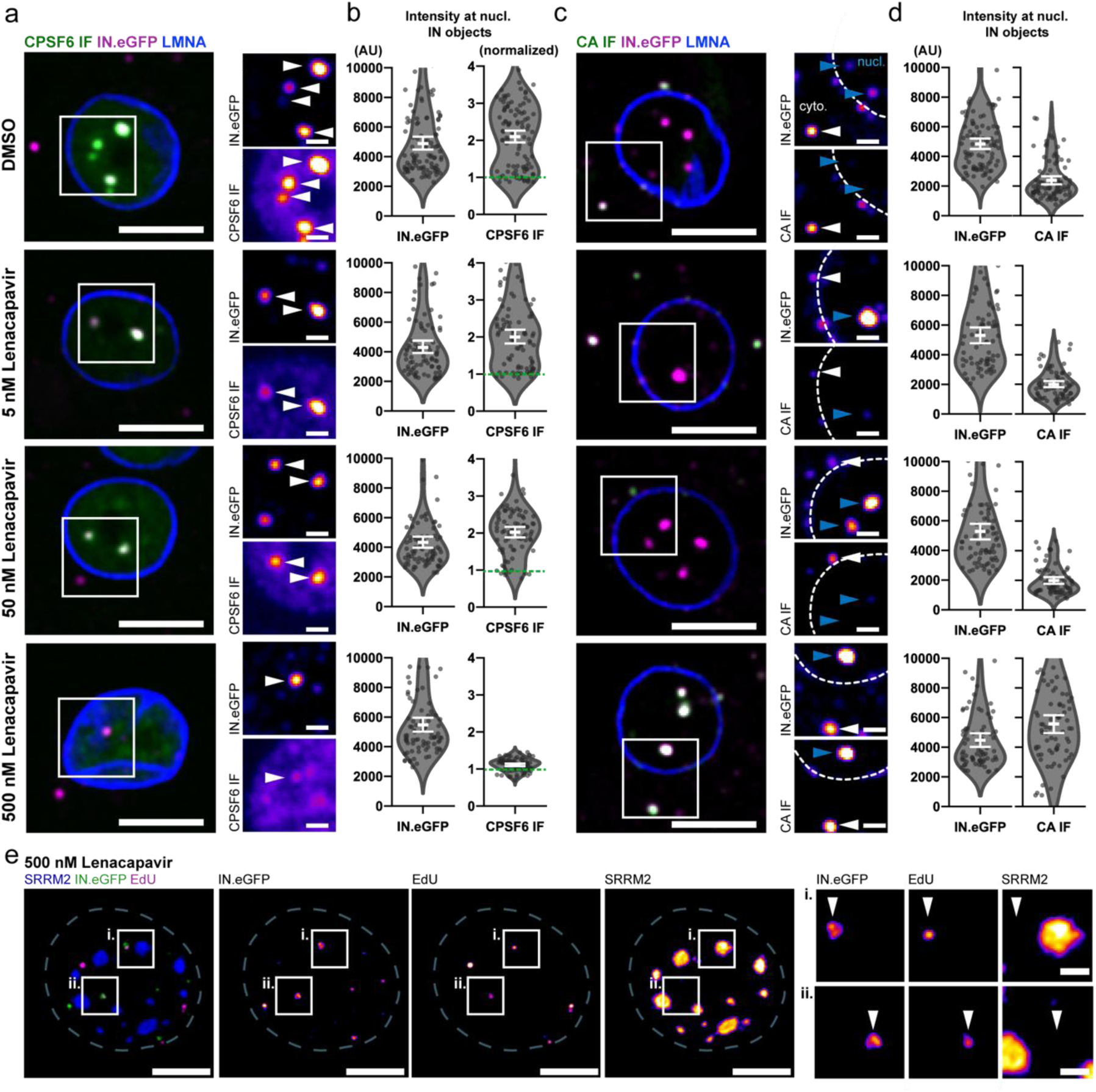
LEN treatment displaces CPSF6 from nuclear capsids, exposes masked CA epitopes and leads to exit of subviral structures from nuclear speckles in MDM. MDM were infected with IN.eGFP labelled VSV-G pseudotyped NNHIV for 72 h prior to addition of indicated concentrations of LEN for 1 h. Cells were fixed and immunostained before SDCM (a-d) or Airyscan (e) 3D imaging. Samples were stained for CPSF6 (a,b), CA (c,d), LMNA (a-d), or SRRM2 (e). The figure shows maximum intensity projections of the central nuclear region (a-d) or a single z plane (e) of one of three (a-d) or two (e) independent experiments. Error bars represent SEM. Scale bars: 5 µm (overviews) and 1 µm (enlargements). **a-d** Displacement of pre-assembled CPSF6 (a,b) and exposure of masked CA epitopes (c,d) by LEN. **a.** White arrowheads indicate nuclear IN.eGFP objects. **c.** White arrowheads indicate cytoplasmic IN.eGFP objects whereas blue arrowheads indicate nuclear IN.eGFP objects. Dotted lines indicate the nuclear boundary. **b,d.** Images were analysed by automated quantification in 3D using custom-made python code as described in materials and methods. CPSF6 signals (b) were normalized to the mean nuclear CPSF6 expression level of the respective cell (green dotted line at y = 1). **e.** MDM were infected in presence of EdU and click labelled prior to immunostaining for SRRM2. IN.eGFP objects displaced from nuclear speckles following treatment with 500 nM LEN for 1 h prior to fixation.

LEN-or PF74-induced removal of CPSF6 from nuclear speckle-associated HIV-1 subviral complexes resulted in their rapid (within 1h) exit from speckles into the adjacent interchromatin space in MDM (Figure 2e and Supplemental Figure 2e,f). This result was similar to the effect of PF74 in model cell lines and T cells, where CPSF6 displacement and exit of HIV-1 complexes were also reported (Francis *et al*, 2020). However, these authors did not observe rapid relocalization of HIV-1 complexes from nuclear speckles in MDM, while this was clearly observed in the current report. Our results indicate a significant concentration difference for LEN-and PF74-induced effects in MDM compared to other cell types and this may also have obscured the effect in the prior study. To evaluate whether tagged IN signals exiting from nuclear speckles correspond to actively reverse transcribing nuclear HIV-1 complexes, we also analyzed infected MDM labeled with EdU and subsequently treated with 500 nM LEN for IN.eGFP and EdU localization with respect to the speckle marker SRRM2 (Figure 2e and Supplemental Figure 2e,f). Following LEN treatment, EdU and IN.eGFP colocalized in the vicinity of nuclear speckles but clearly separated from the SRRM2 signal. We thus conclude that productively reverse transcribing HIV-1 complexes still encased inside the viral capsid exit from nuclear speckles in MDM upon LEN or PF74 induced CPSF6 removal.

### Largely completed nuclear HIV-1 dsDNA can be exposed by treatment of infected MDM with CA targeting drugs

We had previously shown that HIV-1 genomic dsDNA separates from the structural and reporter proteins (IN.eGFP) over time (Müller *et al*, 2021). For this, we made use of the ANCHOR system where a GFP-fused DNA binding protein (OR3) can bind repeats of its cognate binding sequence (ANCH) in the HIV-1 genome, provided that the sequence exists as dsDNA and is accessible to the fusion protein. Since the ANCH repeats were inserted into the viral *env* gene, the system detects only genomes that have been almost completely or completely reverse transcribed into dsDNA. To identify accessible HIV-1 dsDNA genomes and determine their colocalization with and potential alterations of the associated capsid structures, we infected a HeLa-derived cell line stably expressing eGFP.OR3 with NNHIV-ANCH also carrying IN.SNAP as a protein marker. At 23 h p.i., infected cells were treated with 15 µM PF74 or DMSO solvent for 1 h and analyzed for the number of detectable eGFP.OR3 signals in the nucleus and for the colocalization of these signals with IN.SNAP and CA. Most IN.SNAP positive objects did not colocalize with eGFP.OR3 positive objects in DMSO treated control cells (Figure 3a), and eGFP.OR3 positive objects were localized outside of nuclear speckles (Supplementary Figure 3e). The number of eGFP.OR3 positive nuclear objects increased two- to threefold in PF74-treated cells compared to control cells (average of 14 instead of 6 objects per nucleus; Figure 3b,c). Furthermore, the proportion of IN.SNAP/CA signals colocalizing with eGFP.OR3 increased almost threefold from ∼ 25 % measured in control cells to ∼ 70 % in the PF74 treated sample (Figure 3b,d, empty arrowheads). Of note, despite the observed increase of eGFP.OR3 signals inside the nucleus, cytoplasmic IN.SNAP positive objects remained devoid of eGFP.OR3 (Figure 3b, filled arrowheads), consistent with previous reports that reverse transcription is only completed in the nucleus (Dharan *et al*, 2020; Burdick *et al*, 2020; Müller *et al*, 2021).

**Figure 3.**
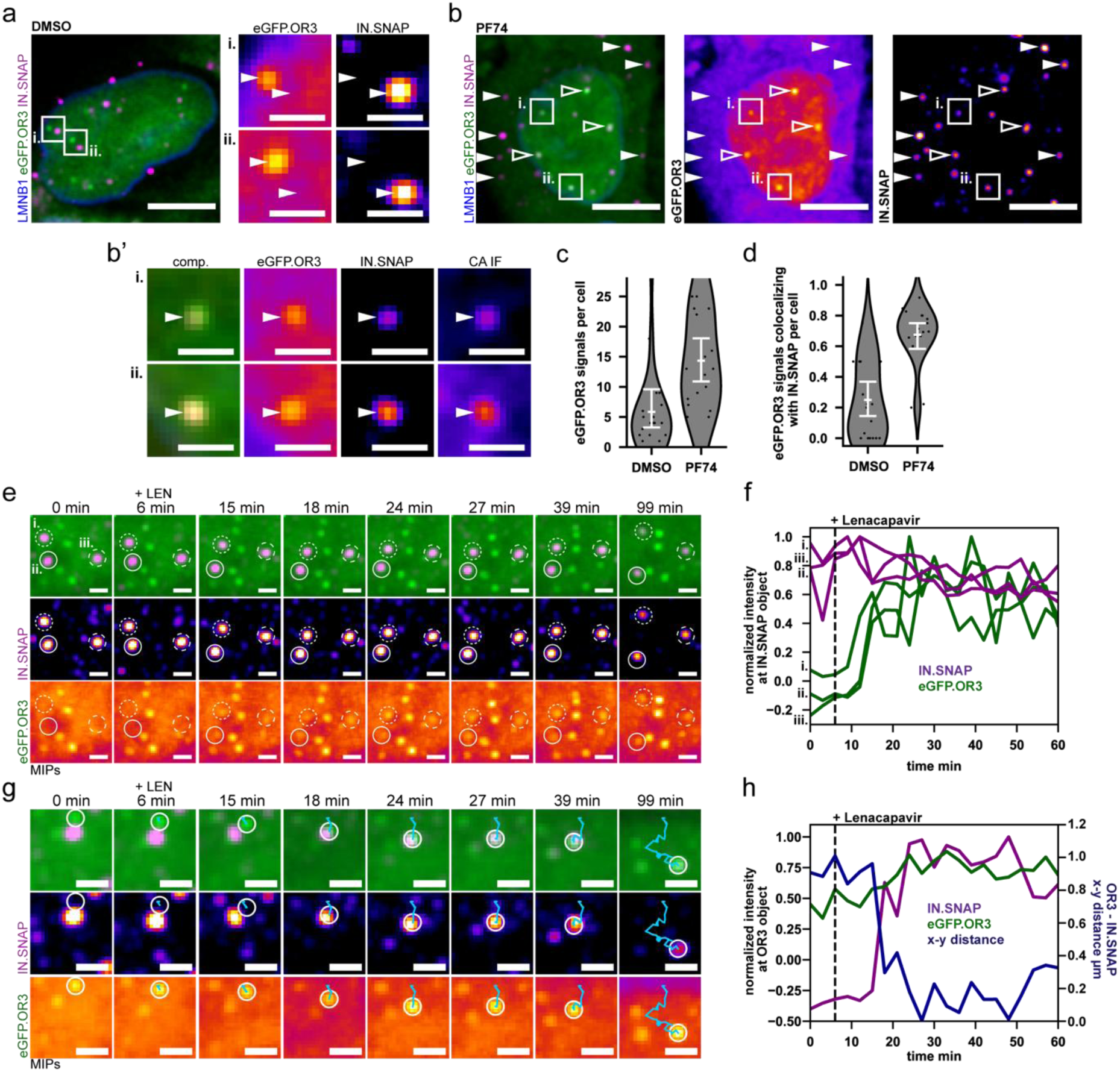
Exposure of nuclear HIV-1 dsDNA (OR3) by treatment with PF74 and LEN. HeLa-based cells stably expressing eGFP.OR3 and eBFP2.LMNB1 were infected using VSV-G pseudotyped NNHIV ANCH IN.SNAP and imaged at 24 h p.i. using 3D confocal spinning disc microscopy. **a.** Representative control cell treated with DMSO for 1 h. Arrowheads in enlargements indicate positions of nuclear eGFP.OR and IN.SNAP punctae, respectively. Scale bars: 5 µm (overview) and 2 µm (enlargements). **b** Cells infected as in (a) and treated with 15 µM PF74 for 1 h before fixation. Arrowheads indicate IN.SNAP punctae in the cytosol (filled) or nucleus (open), respectively. Note that no eGFP.OR3 punctae are detected on IN.SNAP positive objects in the cytosol. Scale bars: 5 µm. **b’.** Enlargements show two exemplary complexes boxed in the nuclear region of (b). Scale bars: 2 µm. **c-d.** Quantification of the total number of nuclear eGFP.OR3 objects per cell (c) and the fraction of eGFP.OR3 signals colocalizing with IN.SNAP objects per cell (d) in DMSO and 15 µM PF74 treated cells. **e-h.** Live imaging of cells using 3D confocal spinning disc microscopy. New eGFP.OR3 signals appear at positions of IN.SNAP objects (e,f) and IN.SNAP objects move to positions of previously present eGFP.OR3 objects (g,h). Recording starts at 22 h p. i. with a time resolution of 3 minutes per frame. 500 nM LEN was added 6 min after start of imaging. **e,g.** Maximum intensity projections (MIPs) of representative nuclear events. See supplementary Movies 2 (e,f) and 3 (g,h). Scale bars: 2 µm **f.** Quantification of signal intensities of the 3 IN objects circled in (e). **h.** Quantification of signal intensities of the OR3 object circled in (g). Distance to the centroid of the IN.SNAP object are presented on the right y-axis. **f,h.** Plotted are mean intensities (diameter 440 nm) normalized with the local background mean intensities (ring diameter 440-1100 nm from centroid) at IN.SNAP (f) or eGFP.OR3 (h) objects in both, the IN.SNAP and the OR3 channels.

Similar results were obtained with LEN (Figure 3e-h and Figure 4a,b and d-f). Live imaging indicated that eGFP.OR3 signals appeared within minutes after addition of LEN at nuclear diffraction limited IN.SNAP signals (which presumably represent clusters of subviral particles based on the CLEM data described above) (Figure 3e-f and Supplementary Movie 2). Subsequently, we observed movement of IN.SNAP/eGFP.OR3 positive clusters to positions of nearby HIV-1 dsDNA signals (eGFP.OR3), which had already been present prior to the start of imaging (Figure 3 g,h and Supplementary Movie 3), consistent with a LEN induced relocation of IN.SNAP objects out of nuclear speckles and into speckle-associated chromatin domains (compare Figure 2e and Supplementary Figure 2e,f). Both effects together explain the observed enhanced colocalization between eGFP.OR3 and IN.SNAP (Figure 3b,d and Figure 4a,b,e). The rapid appearance of new OR3 signals upon drug treatment suggested that nearly complete viral cDNA was already present within the respective subviral complexes, but was still shielded by the viral capsid, the CPSF6 coat or another component of the nuclear speckle.

**Figure 4.**
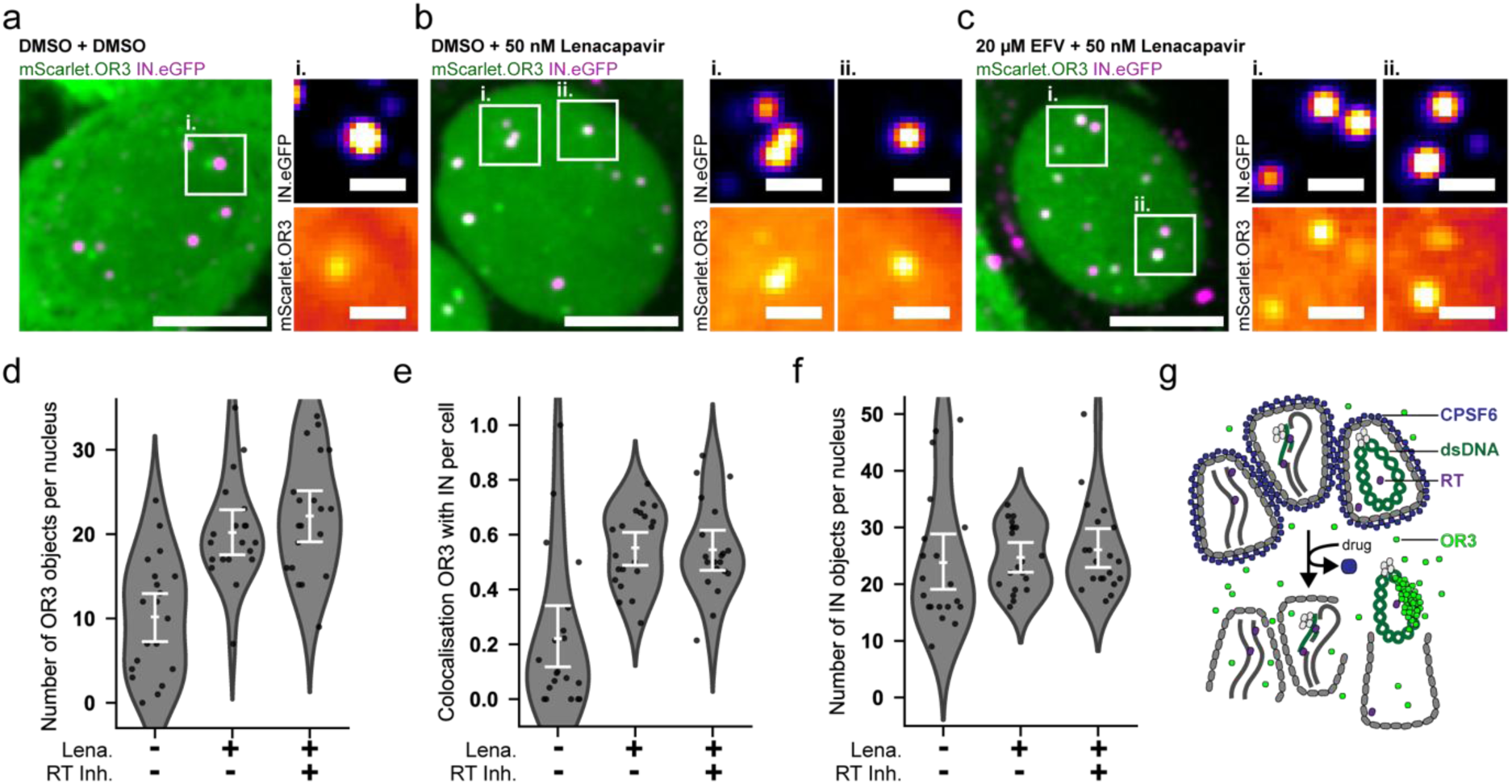
Exposure of additional OR3 signals in the nucleus by LEN treatment occurs independent of ongoing reverse transcription. mScarlet.OR3 expressing TZM-bl cells were infected with VSV-G pseudotyped IN.eGFP labelled NNHIV for 16 h prior to addition of the RT inhibitor EFV or DMSO vehicle and incubated for 45 min, after which 50 nM LEN was added or not for 1 h. **a-c.** Representative examples of cells treated with DMSO/DMSO (a), DMSO/50 nM LEN (b) and 20 µM EFV/50 nM LEN (c). Scale bars: 5 µm (overviews) and 1 µm (enlargements). **d-f.** Quantification of the number of mScarlet.OR3 objects per nucleus (d), the fraction of mScarlet.OR3 signals colocalizing with IN.eGFP per nucleus (e), and the number of IN.eGFP objects per nucleus (f). Error bars represent 95 % CI and points represent single cells. g. Scheme of LEN/PF74 induced exposure of dsDNA.

The appearance of new OR3 signals colocalizing with HIV-1 subviral complexes could also be due to previously stalled reverse transcription complexes resuming reverse transcription to complete dsDNA upon opening of the capsid. To evaluate this possibility, the reverse transcriptase inhibitor Efavirenz (EFV) was added 45 min before addition of LEN to cells infected for 16 h (Figure 4). This treatment blocks any further reverse transcription immediately before and during subsequent LEN treatment. The average number of mScarlet.OR3 foci per nucleus (Figure 4d) and the colocalization between IN.eGFP and mScarlet.OR3 were again two-to threefold higher in LEN treated cells compared to the DMSO control (Figure 4e), independent of the absence or presence of EFV. The average number of IN.eGFP positive objects per nucleus was not affected by these treatments (Figure 4f). These results showed that the mScarlet.OR3 signals newly detected upon LEN addition represented viral cDNA molecules that had already undergone reverse transcription to complete or almost complete dsDNA, but were not accessible to binding by the mScarlet.OR3 marker before LEN-induced removal of the CPSF6 coat (Figure 4g).

### LEN treatment alters capsid morphology in infected cells

Drug-induced accessibility of pre-synthesized nuclear HIV-1 dsDNA indicated that LEN or PF74 binding may affect capsid structure and integrity besides removing its CPSF6 coat, in accordance with prior *in vitro* experiments (Faysal *et al*, 2024). In order to morphologically characterize potential drug induced changes on nuclear HIV-1 capsids, we employed CLEM-ET. We infected mScarlet.OR3 expressing cells with IN.SNAP.SiR labelled NNHIV for 24 h and treated the cells with 500 nM LEN, 15 µM PF74 or DMSO for 1 h before cryoimmobilization by high-pressure freezing, freeze-substitution and plastic embedding. Electron tomograms were recorded at positions correlated to either nuclear IN.SNAP.SiR objects or mScarlet.OR3 objects.

As observed previously (Müller *et al*, 2021), tomograms correlated to nuclear IN.SNAP positive positions (independent of additional mScarlet.OR3 signal presence) in control cells consistently revealed clusters of capsid-like objects often containing electron-dense material inside (Figure 5a panels i.), whereas tomograms recorded at mScarlet.OR3 positions lacking IN.SNAP did not show a defined structure but rather a density distribution resembling surrounding chromatin (Figure 5a panels ii.). In PF74-treated cells, tomograms were recorded at IN.SNAP.SiR positive positions that often were adjacent to, but not directly co-localizing with mScarlet.OR3. We observed structures closely resembling conical HIV-1 capsids that often contained electron-dense material inside. Some of these structures exhibited thin, electron-dense protrusions emanating from the narrow end of the cone-shaped capsid (Figure 5b, cyan arrows). The same region also revealed broken capsid-like remnants (Figure 5b). The addition of 500 nM LEN had a similar effect, again showing dense clusters of cone-shaped structures with electron-dense material inside and resembling HIV-1 capsids (Figure 5c,d). The arrangement of HIV-1 complexes appeared to be more condensed and electron-dense compared to the DMSO control and to previous results from untreated cells (Zila *et al*, 2021; Müller *et al*, 2021; Schifferdecker *et al*, 2022). Given that native capsids isolated from HIV-1 particles aggregate rapidly into dense clusters (Welker *et al*, 2000), we attribute this observation to the removal of the CPSF6 coat from nuclear HIV-1 capsids. Furthermore, striking electron-dense protrusions were frequently observed, always emanating from the narrow end of cones (Figure 5c cyan arrowheads; figure 5d, white arrowheads). In addition to these characteristic structures, 3D rendering of tomographic reconstructions revealed abundant broken and apparently connected empty lattices (Figure 5d red stars), seemingly oriented with their narrow ends towards clusters of structures (Figure 5d, Supplementary Movie 4).

**Figure 5.**
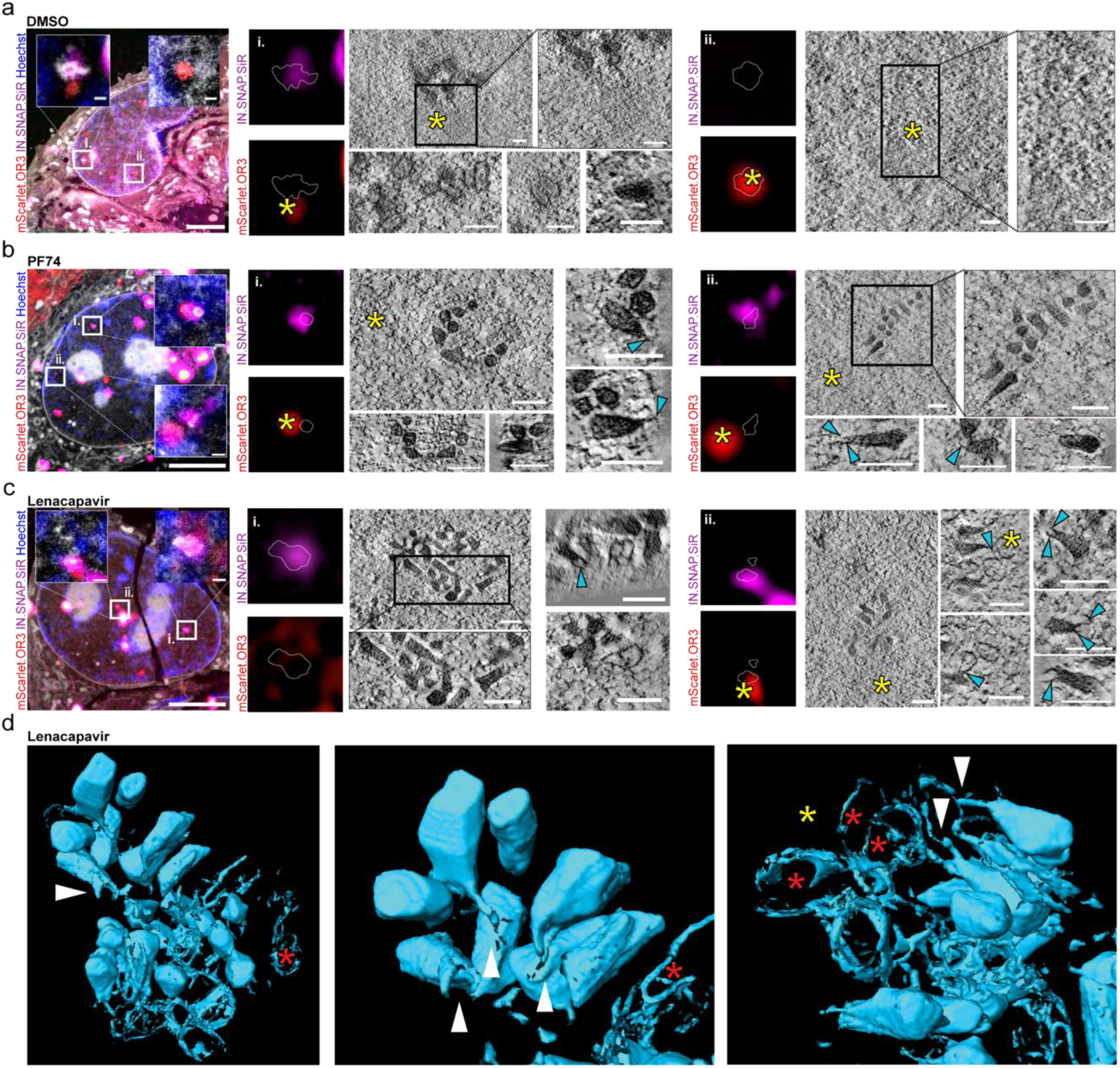
Correlative light and electron microscopy with electron tomography (CLEM-ET) analyses of capsid alterations following LEN/PF74 treatment. mScarlet.OR3 expressing HeLa-based TZM-bl cells infected with VSV-G pseudotyped IN.SNAP.SiR labelled NNHIV were treated at 24 h p.i. with DMSO (a), 15 µM PF74 (b) or 500 nM LEN (c,d) for 1 h prior to cryoimmobilization, freeze-substitution and further processing for CLEM. Electron tomograms were acquired at positions correlated to IN.SNAP.SiR and/or mScarlet.OR3 signals. **a-c.** Left panels show CLEM overlays of the 250 nm thin sections with the inverted EM image, positive for mScarlet.OR3 (red) and/or IN.SNAP.SiR (magenta), post-stained with Hoechst (blue), and decorated with multifluorescent fiducials for correlation (all channels, white). Slices of individual tomograms correlated to the indicated positions are presented. Cyan arrowheads indicate protrusions from narrow ends of conical capsid objects. Scale bars: 2.5 µm (overlay) 100 nm (tomograms). **a.** Tomograms of DMSO treated cells correlated to an IN.SNAP.SiR signal with an mScarlet.OR3 signal in close proximity (yellow star, position i.) and correlated to an mScarlet.OR3 signal (yellow star) lacking IN.SNAP.SiR (position ii.). **b.** PF74 treated cells correlated to IN.SNAP.SiR signals associated with mScarlet.OR3 signals (yellow stars, both positions). **c.** LEN treated cells with presented tomograms correlated to IN.SNAP.SiR positions lacking mScarlet.OR3 (position i.) or overlapping with mScarlet.OR3 signal (yellow star, position ii.) **d.** 3D rendering of segmented electron densities of tomogram shown in (c, position ii.). Red stars indicate empty/fused lattices and white arrowheads point to protrusions at the narrow ends of conical capsid objects. See Supplementary Movie 4.

## Discussion

While many recent studies strongly enhanced our understanding of the early phase of HIV-1 replication, the processes of intranuclear trafficking, genome uncoating, and how they relate to integration are still incompletely understood. Here we showed that morphologically intact HIV-1 capsids with interior nucleic acid density as well as broken and empty capsid-like particles clustered in nuclear speckles of post-mitotic primary MDM in a CPSF6-dependent way, as previously observed in various tissue culture adapted cell lines (Müller *et al*, 2021; Schifferdecker *et al*, 2022; Ay *et al*, 2025). Treatment of cells containing nuclear HIV-1 complexes with capsid-targeting drugs led to CPSF6 removal, exit from nuclear speckles and accessibility of pre-synthesized full-length or near full-length HIV-1 dsDNA, indicating genome uncoating. The observation of broken capsids without interior density is consistent with prior results in different cell lines that also reported incomplete capsid lattice remnants inside the nucleus of HIV-1 infected cells (Müller *et al*, 2021; Ay *et al*, 2025). These results indicate physical breakage of the capsid as general mechanism of HIV-1 genome uncoating rather than disassembly of the CA lattice. Other studies suggesting rapid disassembly of the capsid lattice (Burdick *et al*, 2020; Li *et al*, 2021) applied substoichiometric amounts of a GFP-tagged CA fusion protein as a marker incorporated into HIV-1 capsids rather than EM detection of the actual structures. This fusion protein may conceivably be preferentially lost from the broken capsid lattice, thus mimicking capsid disassembly.

HIV-1 capsids associated with CPSF6, clustered within the center of nuclear speckles of MDM, and contained *de novo* synthesized viral DNA. Super-resolution STED analysis of the major nuclear speckle component SRRM2 indicated distinct speckle subdomains, consistent with findings from HeLa, HEK293T, and WI-38 (human lung fibroblast) cells obtained using structured illumination microscopy (SIM) (Fei *et al*, 2017; Zhang *et al*, 2024) and stochastic optical reconstruction microscopy (STORM) (Zhang *et al*, 2024). SRRM2 and the second major nuclear speckle component SON inhabit distinct immiscible phases in nuclear speckles, each enriched with different speckle components (Zhang *et al*, 2024). These substructures likely represent clusters of ∼ 40 nm ordered assemblies driven by microphase separation of block copolymer-like splicing factors (Shinn *et al*, 2025). Interestingly, we observed that capsid clusters localized to the subdomains lacking SRRM2. We hypothesize that not only the immersion of capsids into the phase-separated speckle environment (Xu *et al*, 2022), but also this specific subdomain localization is driven by the biophysical properties of the multimeric CPSF6 coat encasing nuclear capsids. Our live imaging and CLEM data showed that removing the CPSF6 layer by LEN or PF74 treatment forced the clustered capsids to move out of nuclear speckles in a concerted fashion, remaining densely clustered in the adjacent nucleoplasm. Of note, these structures often followed the path of previously uncoated viral dsDNA, which had dissociated from the viral structures inside nuclear speckles before drug treatment. While this may simply reflect the dense 3D chromatin architecture constraining possible trajectories, it is also conceivable that specific interactions between the exposed dsDNA and the remaining subviral complex influence these dynamics.

Treatment of infected cells carrying nuclear HIV-1 complexes with LEN or PF74 induced exposure of full length or nearly full-length viral dsDNA to be recognized by a 66 kDa fusion protein binding a specific DNA sequence in the dsDNA genome. This effect was independent of ongoing reverse transcription, indicating that complete or almost complete dsDNA of HIV-1 can be synthesized within an apparently intact HIV-1 capsid that excludes access of the 66 kDa fluorescent fusion protein to the capsid interior. This observation is consistent with the ability of HIV-1 to evade activation of innate immune responses (Cingöz & Goff, 2019) unless cGAS-mediated sensing is induced by PF74 (Sumner *et al*, 2020; Ay *et al*, 2025) or LEN (Scott *et al*, 2025). Our findings suggest that dsDNA synthesis of the viral genome may be important for capsid breakage and genome uncoating, but does not appear to be sufficient for these processes. ANCHOR detection in our system requires the dsDNA to be completed to at least 7.3 kbp. This DNA length is well above the critical limit of ∼ 6 kbp dsDNA for efficient HIV-1 genome uncoating reported in a previous study (Burdick *et al*, 2024). We hypothesize, therefore, that additional factors besides complete dsDNA synthesis are important to reveal the HIV-1 genome to the nuclear environment. Given that capsid-targeting drugs remove the CPSF6 coat and induce capsid breakage, we cannot currently distinguish which of these mechanisms is most relevant for the observed phenotype. Drug induced displacement of the CPSF6 coat could make the capsid more sensible to breakage by dsDNA or could expose a previously broken capsid lattice that had been stabilized by the outer CPSF6 coat. Alternatively, genome exposure may depend on the direct effect of PF74 or LEN binding on capsid lattice stability that has been demonstrated *in vitro* (Faysal *et al*, 2024). Conceivably, capsid integrity may become compromised already upon capsid passage through the narrow NPC channel. This was not apparent in a recent cryo-electron tomography study, but complete tracing of the lattice of NPC-associated and nuclear HIV-1 capsids was not possible due to low number of events and dense surrounding environment (Kreysing *et al*, 2025). In the described scenario, CPSF6 coating of the capsid as it exits the NPC channel would shield and stabilize the subviral structure, and removal of CPSF6 may then be sufficient for breaking the lattice by the forces of internal dsDNA. We can currently not distinguish which of these effects, or a combination thereof, are critical for drug induced DNA exposure in our experiments. Independent of the precise mechanism, our results together with those of Burdick *et al*. (Burdick *et al*, 2024), indicate that synthesis of long dsDNA and factor(s) affecting the integrity of the subviral structure in nuclear speckles may act together to induce efficient capsid opening and genome uncoating and may also ensure that this occurs at the optimal position for genome integration into transcriptionally active chromatin. This may not be an all-or-none phenomenon, but rather influence the kinetics and efficiency of uncoating. Furthermore, these processes may be affected by the availability of host factors in different cell types, consistent with the observation of different LEN concentrations required for CPSF6 removal and DNA exposure depending on the cell type.

LEN treatment resulted in a clear change of HIV-1 capsid morphology within the nucleus, frequently inducing bifurcated linear protrusions emanating from the narrow ends of capsid cones. These protrusions could represent structurally altered parts of the capsid lattice, and/or the (chromatinized) viral dsDNA complex emanating from an opening in the capsid. We did not observe direct colocalization of the HIV-1 dsDNA-binding fusion protein with IN.SNAP.SiR positive positions, but this may have been due to fluorescence-intensity losses by CLEM sample preparation, while brighter mScarlet.OR3 signals that existed prior to drug treatment were retained. Clusters of capsids displaying such electron-dense protrusions appeared in close proximity to networks of empty and flattened lattice structures possibly connected to each other, consistent with the observation that LEN and related compounds both disrupt the integrity of the mature HIV-1 capsid and at the same time hyperstabilize the hexameric CA lattice (Faysal *et al*, 2024; Li *et al*, 2025). Given that LEN binding is incompatible with CA pentamers (Huang *et al*, 2025), we speculate that local conversion of pentamers may lead to disruption and opening of the capsid structure. Pentamer density is highest at the narrow end of the cone, which can explain the preferential deformation and opening of capsids in this region allowing release and chromatinization of the viral genome from the capsid interior upon drug treatment. Drug-induced stabilization of the remaining hexameric capsid lattice can also explain the high prevalence of flattened or apparently fused lattice remnants in our study. Mechanistically this might be achieved by binding of PF74 and LEN to the Thr-Val-Gly-Gly motif within CA, a region which mediates a structural switch between a pentamer- and hexamer-favoring conformation (Schirra *et al*, 2023; Stacey *et al*, 2023). The altered capsid morphology we observed in the nuclei of infected cells is consistent with very recent negative-stain electron micrographs of isolated capsids treated with LEN *in vitro* (Li *et al*, 2025). However, this does not necessarily reflect the regular process of uncoating in the absence of drugs, since endogenous reverse transcription in isolated native HIV-1 capsids *in vitro* induced lattice breakage and polynucleotide loop emanation at the broad end and long side of the capsid as well (Christensen *et al*, 2020).

In summary, our results point to additional factor(s) involved in capsid breakage and genome release besides synthesis of a sufficiently long dsDNA contributing the physical force. These may include factors destabilizing the CPSF6 coat from speckle-associated capsids and/or factors destabilizing the capsid lattice. Once the capsid is broken, rapid chromatinization of exposed dsDNA loops (Christensen *et al*, 2020) via histones (Geis & Goff, 2019; Wang *et al*, 2016) or binding of other proteins to exposed DNA loops may facilitate genome exit from broken capsid remnants. Understanding the detailed mechanisms and factors involved in nuclear HIV-1 genome uncoating and subsequent integration can also help to better understand latency induction and maintenance, with the aim to devise efficient strategies to purge the latent reservoir in infected individuals. Our results also shed further light on the mechanism(s) of action of the potent anti-HIV drug LEN.

## Materials and Methods

### Cell culture

Hela-based TZM-bl cells (Wei et al, 2002) and human embryonic kidney 293T cells (HEK293T) (Pear et al, 1993) were cultured in Dulbecco’s Modified Eagle’s Medium (DMEM) supplemented with 10 % fetal bovine serum (FBS), 100 U ml^1^ penicillin, and 100 mg ml^1^ streptomycin (PAN Biotech, Germany). eGFP.O3, mScarlet.OR3 and eBFP2.LMNB1 expressing TZM-bl cells have been described previously (Müller *et al*, 2021). Cell lines were tested for mycoplasma contamination (MycoAlert mycoplasma detection kit, Lonza Rockland, USA) and authenticated by STR profiling (Eurofins Genomics, Germany). MDM were isolated from buffy coats of healthy human blood donors and cultured in RPMI 1640 (Thermo Fisher Scientific) containing 10 % heat-inactivated FBS, 100 U ml^1^ penicillin, 100 mg ml^1^ streptomycin and 5 % human AB serum (Sigma Aldrich) as described previously (Bejarano et al, 2019).

### Virus stock production

293T cells were seeded into nine T175 dishes (per ultracentrifuge rotor) to reach 50-70 % confluency the next day. Cells were then transfected with proviral plasmid NNHIV env(stop) ANCH (Müller *et al*, 2021), IN.SNAP or IN.eGFP expression plasmid (Müller *et al*, 2021), and pCMV-VSV-G (Addgene plasmid #8454, a gift from from Bob Weinberg) at a ratio of 7.7: 1.3: 1.0 µg using calcium phosphate (70 µg total DNA per T175 dish). Medium was changed after 4-8 hr and cells were incubated at 37 °C for 2 days prior to harvesting the supernatant. Centrifugation at 300 g was performed for 5 minutes before filtration through 0.45 µm mixed cellulose ester (MCE) filters. Filtered supernatant was overlayed onto 20 % (w/w) sucrose and centrifuged at 107 000 g for 2 h at 4 °C. Viral particles were resuspended in 75 µl PBS containing 10 % FCS and 10 mM HEPES (pH 7.5), pooled and frozen at - 80°C. Viral stocks were quantified using the SYBR Green based Product Enhanced Reverse Transcription assay (SG-PERT) as described previously (Pizzato et al, 2009).

### Virus labelling, infection of cells, and drug treatments

3,33 x 10^3^ TZM-bl cells or 5 x 10³ MDM per well were seeded into 15 well µ-Slides (Ibidi, Germany) in 50 µl medium and incubated over night at 37 °C and 5 % CO2. BG-SiR (SNAP-SiR, New England Biolabs) was prediluted to 20 µM in medium and added 1:10 to concentrated particles for a final concentration of 2 µM and incubated for 30 minutes at 37 °C. Cells were infected using 30 µU (TZM-bl) or 60 µU (MDM) RT activity of IN.eGFP or IN.SNAP labelled VSV-G pseudotyped NNHIV env(stop) ANCH. Cells were drug treated and fixed at indicated timepoints. 20 µM Efavirenz (EFV, Sigma-Alldrich) was added 45 min prior to indicated concentrations of Lenacapavir (LEN or GS-6207, provided by Mamuka Kvaratskhelia, University of Colorado) or 15 µM PF74 (Sigma-Alldrich). LEN and PF74 were added 1 h prior to washing the cells with PBS and fixation with 4 % paraformaldehyde (PFA). EdU (Thermo Fisher Scientific) was added at the time of infection at 10 µM.

### Immunofluorescence staining and EdU click labeling

Fixed cells were permeabilized by 0.5 % Triton X-100 for 10 minutes, followed by washing 3 times with 3 % bovine serum albumin (BSA) in PBS and blocking for 1 h at room temperature. In case of EdU click labeling, cells were treated with Click-iT EdU Alexa Fluor 647 Imaging kit (Thermo Fisher Scientific, USA) according to the manufacturers’ instructions. Primary antibodies (table 1) were diluted in 0.5 % BSA in PBS and incubated for 1 h at room temperature. After 3 washing steps with 3 % BSA in PBS, secondary antibodies (anti-rabbit or anti-mouse goat polyclonal labelled with Alexa Fluor 405, 488, 568, or 647 (Thermo Fisher Scientific) for confocal and Airyscan microscopy, or labelled with STAR RED (Abberior) or Atto 594 (Sigma-Aldrich) for STED microscopy) were diluted 1:1000 (Alexa Fluor labelled antibodies) or 1:500 (STAR RED and Atto 594 labelled antibodies) with 0.5 % BSA in PBS and added to the cells. After 1 h incubation at room temperature in the dark, cells were washed 3 times with PBS and stored at 4 °C in the dark until imaging.

**Table 1.** primary antibodies

| Target | Source | Supplier | Cat no. or reference | Dilution |
| --- | --- | --- | --- | --- |
| anti-hLamin A/C | mouse monoclonal | Santa Cruz Biotech. | Cat#:sc-7292 | 1:100 |
| anti-hCPSF6 | rabbit polyclonal | Atlas Antibodies | Cat#:HPA039973 | 1:500 |
| Anti-CA antiserum | rabbit polyclonal | in-house | (Bejarano *et al*, 2019) | 1:1000 |
| anti-SON antibody | rabbit polyclonal | Sigma-Aldrich | Cat#: HPA023535 | 1:800 |
| anti-SRRM2 (SC35) | mouse monoclonal | Abcam | Cat#: ab11826 | 1:500 |

### Imaging

Confocal spinning disc imaging was performed using an inverted Perkin Elmer Ultra VIEW VoX 3D spinning disk confocal microscope (Perkin Elmer, United States) with a 60x oil immersion objective (NA 1.49; Perkin Elmer) and a pixel size of 0.22 µm. 3D stacks of 10–30 randomly chosen positions were automatically recorded with a z-spacing of 0.5 µm. For live cell imaging, medium was changed to imaging medium containing FluoroBrite DMEM (Thermo Fisher Scientific), 10% FCS, 4 mM GlutaMAX (Gibco Life Technologies), 2 mM sodium pyruvate (Gibco Life Technologies), 20 mM HEPES pH 7.4, 100 U ml^-1^ Penicillin and 100 µg ml^-1^ Streptomycin (PAN-Biotech) and cells were imaged inside a humid incubation chamber set to 37 °C and 5 % CO2 with a time interval of 3 minutes per stack.

STED imaging was performed using a STED system (Abberior Instruments GmbH, Germany) with a 775 nm depletion laser and 100x oil immersion objective (NA 1.4; Olympus UPlanSApo). Images were acquired in the 590 and 640 laser lines with a nominal STED power set to 80 % of the maximal power of 3 W, 20 µs pixel dwell time and 15 nm pixel size. STED images were deconvolved using the Richardson-Lucy algorithm within the Imspector software (Abberior Instruments GmbH).

Airyscan point laser scanning confocal microscopy was performed on a Zeiss LSM900 microscope (Carl Zeiss Microscopy, Germany) equipped with the Airyscan detector, using an oil immersion Plan-Apochromat 63x objective (NA 1,4/Zeiss) in the Airyscan super resolution mode. Multichannel images were acquired sequentially in the stack scanning mode using 405 nm, 488 nm and 640 nm diode lasers for fluorophores in the blue, green and far-red spectrum, respectively. Emission detection was configured using variable dichroic mirrors to be 400-490 for blue fluorophore detection, 490-580 for green fluorophore detection and 620-700 for far red fluorophore detection. Airyscan detectors were used with the gain adjusted between 700 and 900 V; offset was not adjusted (0%). Sampling was system-optimized for Airyscan super-resolution imaging (approx. 50 nm in xy axis and 130 nm in z axis), acquisition was performed bidirectionally with a pixel dwell time between 0.7 and 1.2 μs. Subsequently, ZEN Blue 3.1 software was used for 3D Airyscan processing with automatically determined default Airyscan Filtering (AF) strength.

### Image analyses and data visualization

Presentation of images. For visualization purposes, confocal and Airyscan images were filtered using a 1 pixel mean filter using Fiji/ImageJ (Schindelin *et al*, 2012) to reduce noise. The standard ‘Fire’ lookup table within Fiji was used for presenting single channel images in figures. Linear unmixing was performed in case of fluorophore crosstalk.

Quantification of images. An automated processing pipeline was established using a combination of in-house python v3.11.5 code and Ilastik v1.3.3 (Berg *et al*, 2019). A pixel and object classifier was trained using Ilastik on raw 3D stacks of images to segment the Lamin signal and IN.SNAP spots. Object identities were exported and loaded into a python script that segmented the nucleoplasm by mathematically filling the nuclear interior and eroding the mask to exclude the nuclear envelope. IN.SNAP objects were then evaluated based on their pixel location. Only objects with all pixels falling within the nucleoplasmic mask were labelled as nuclear. Mean pixel intensities of these nuclear objects were then collected and plotted. CPSF6 signals were normalized to the mean background expression level of the entire nucleoplasm. For quantification of OR3 signals per nucleus, a semi-automated approach using Icy (de Chaumont *et al*, 2012) was performed. Using the spot detector of Icy, OR3 and IN.SNAP objects of single nuclei were segmented in 3D. All detected objects were subsequently manually corrected and the total number of objects per nucleus was recorded.

Tracking and quantification of live movies was performed using the Trackmate plugin of Fiji/ImageJ and the Track Manager of Icy. Movies were filtered using a mean filter size of 1 pixel and maximum intensity projections were generated using Fiji/ImageJ. Drift in x/y was corrected using the registration function correct 3D drift of Fiji/ImageJ and IN.SNAP/OR3 objects were detected using a LoG (Laplacian of Gaussian) filter and tracked using the simple LAP tracker or manually. Tracks were exported to Icy and with the Track Manager plugin the local background corrected mean intensities were calculated using the intensity profiler track processor with a disk diameter of 2 pixels (440 nm) and a local background ring diameter of 2-5 pixels (440-1100 nm). Distances between tracks were calculated using the distance profiler track processor. Intensities were normalized to the highest value and plotted using the statistical visualization toolbox seaborn v0.13.0 (Waskom, 2021) in python.

### CLEM sample preparation and electron tomography

1.2 × 10^5^ TZM-bl mScarlet.OR3 cells or 4 × 10^4^ MDM were seeded on 3 mm sapphire discs in a 35 mm glass-bottom dish (MatTek, USA), on the next day infected with VSV-G pseudotyped IN.SNAP.SiR labeled NNHIV ANCH (30 µU RT/cell for TZM-bl cells and 60 µU RT/cell for MDM) and incubated for 24 hr at 37°C. Cells were treated with 500 nM LEN, 15 µM PF74, or DMSO vehicle for 1 h prior to cryo-immobilization and electron tomography as described previously (Müller *et al*, 2021).

## Acknowledgements

We are grateful to M. Kvaratskhelia, University of Colorado for a kind gift of LEN. The ANCHOR system is developed by and available from NeoVirTech (France, http://www.neovirtech.com). We acknowledge the microscopy support from the Infectious Diseases Imaging Platform (IDIP) of the Center for Integrative Infectious Disease Research, Heidelberg. This work was funded by the Deutsche Forschungsgemeinschaft (DFG, German Research Foundation) – Projektnummer 240245660 – SFB 1129 project 5 (H-GK), project 6 (BM), project 4 (USS), and by the TTU HIV in the DZIF (VL, H-GK).

## Supplementary Figures

**Supplementary Figure 1.**
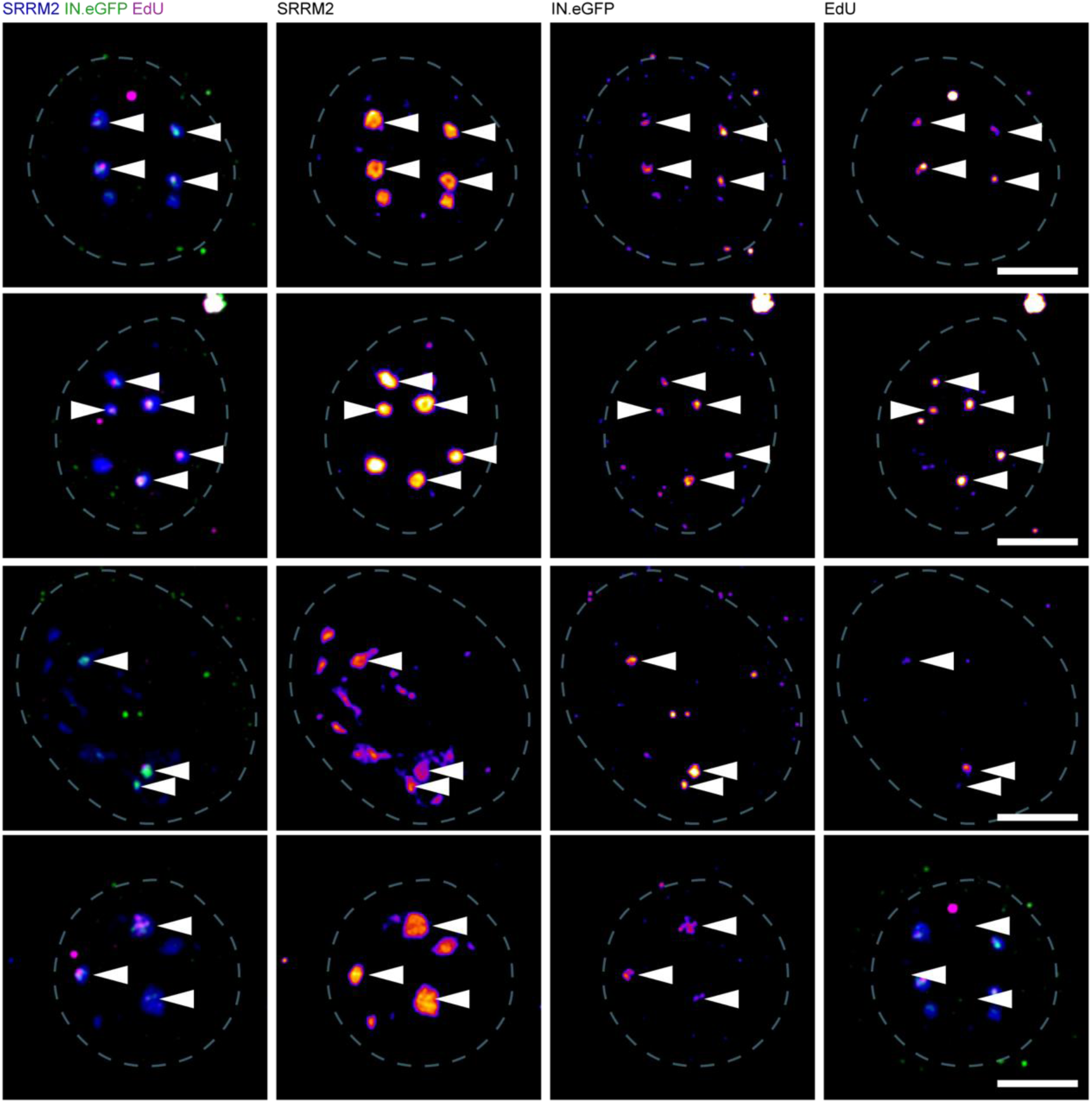
Additional examples of subviral complexes in nuclear speckles of primary MDM. Super-resolution analysis of HIV-1 cDNA within nuclear speckles of monocyte-derived macrophages (MDM) showing EdU (magenta) and IN.eGFP signals (green) in the center of SRRM2 condensates (blue). Shown are four additional maximum intensity projection of MDM nuclei (white dashed line) infected for 72 h with VSV-G pseudotyped NNHIV in presence of EdU followed by fixation, EdU click labeling and immunofluorescence staining using an antibody against SRRM2 (SC35). Samples were imaged using Airyscan microscopy. Scale bars: 5 µm.

**Supplementary Figure 2.**
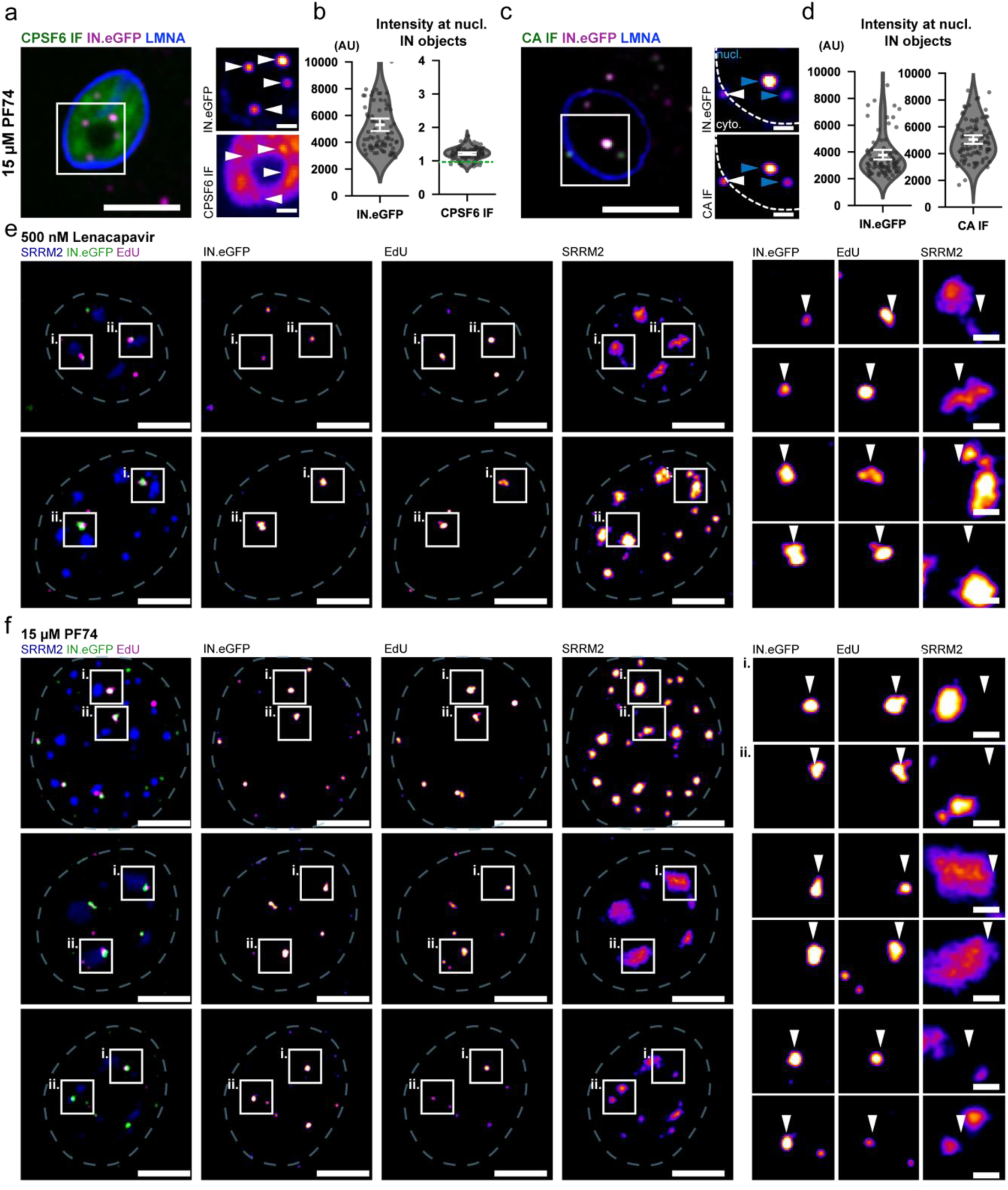
Effect of PF74 on CA and CPSF6 signals in MDM and additional examples of LEN and PF74 induced subviral particle exit from nuclear speckles. **a-d.** MDM were infected using IN.eGFP labelled VSV-G pseudotyped NNHIV IN.SNAP for 72 h before addition of indicated concentrations of PF74 or LEN for 1 h. Cells were fixed and immunostained before 3D SDCM imaging (a-d) or 3D Airyscan imaging (e,f). Samples were stained for CPSF6 (a,b), CA (c,d) or SRRM2 (e,f). Error bars represent SEM. Scale bars: 5 µm (overviews) and 1 µm (enlargements), maximum intensity projection are shown. **a-d** Stripping of pre-assembled CPSF6 (a,b) and exposure of masked CA epitopes (c,d) by 15 µm PF74. **c.** White arrowheads indicate cytoplasmic IN.eGFP objects whereas blue arrowheads indicate nuclear IN.eGFP objects. Dotted lines indicate nuclear boundary. **b,d.** Images were analysed by automated quantification using custom-made python code as described in materials and methods. CPSF6 signals (b) were normalized to the mean nuclear CPSF6 expression level of the respective cell (green dotted line at y = 1). **e,f.** Displacement of IN.eGFP objects from nuclear speckles. Cells were infected in presence of 10 µm EdU and click labelled prior to immunofluorescence staining. Shown are 2 more representative cell nuclei treated with 500 nM LEN for 1 h (e) and three nuclei treated with 15 µM PF74 for 1 h (f).

**Supplementary Figure 3.**
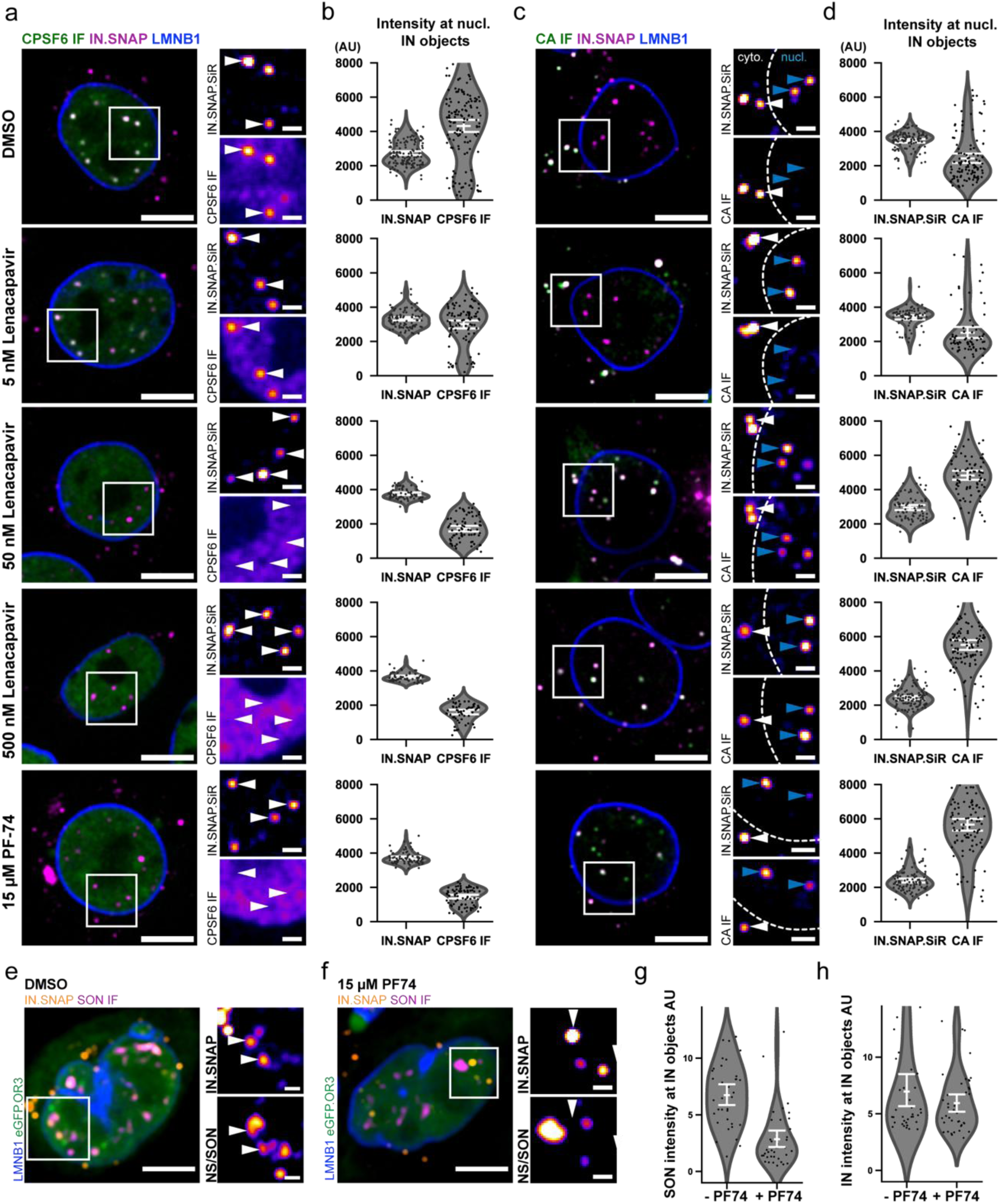
Lenacapavir and PF74 strip CPSF6 from nuclear capsids, concomitantly expose masked CA epitopes, and lead to capsid exit from nuclear speckles of HeLa-based TZM-bl cells. Cells were infected using IN.SNAP labelled VSV-G pseudotyped NNHIV for 24 h before addition of indicated amounts of Lenacapavir, PF74 or DMSO for 1 h. Cells were fixed and immunostained before 3D confocal spinning disc imaging. Samples were stained for CPSF6 (a,b), CA (c,d) or SON (e-g). Shown is one of three independent experiments. Error bars represent SEM. Scale bars: 5 µm (overviews) and 1 µm (enlargements). **a-d** Stripping of pre-assembled CPSF6 (a,b) and concomitant exposure of masked CA epitopes (c,d) by Lenacapavir and PF74. **c.** White arrowheads indicate cytoplasmic IN.eGFP objects whereas blue arrowheads indicate nuclear IN.eGFP objects. **b,d.** Images were analysed by automated quantification using custom-made python code. Nuclear 3D IN.eGFP objects were segmented and mean intensities in the respective channels quantified. **e-h.** Displacement of IN.SNAP objects from nuclear speckles. Shown are representative cells treated with DMSO (e) or 15 µM PF74 (f) for 1 h. **g,h.** Quantification of SON (g) or IN.SNAP (h) mean intensities shown in (e,f) at nuclear 3D IN.SNAP objects in presence and absence of 15 µM PF74.

